# A general route to volumetric segmentation under annotation scarcity

**DOI:** 10.64898/2026.09.16.752170

**Authors:** Samhita Radhakrishnan, Saurabh Mathur, Aleksandr Aleksandrov, Hugo Schweke, Gerardine Silvano Gargano, Emmanuel Levy

**Author notes:** Denotes co-first authorship (Samhita Radhakrishnan, Saurabh Mathur).

## Abstract

Volumetric cell segmentation is limited by the scarcity of labelled data, because manual curation of three-dimensional datasets is prohibitively laborious. We find that this gap can be bridged with volumes in which synthetic 3D objects mirror the properties of real cells and of image formation, notably the axial spread of the point-spread function. Such volumes provide unlimited labelled training data, and we show that a model trained on them produces accurate segmentations of real biological data.

## Main text

Fluorescence microscopy is the workhorse of quantitative cell biology, with advances in optics and automation enabling high-throughput imaging of cells, tissues, and organisms across many conditions (1–3). Extracting quantitative information from these images requires robust segmentation, for which deep learning is driving a performance revolution. Segmentation in two dimensions (2D) is largely solved by generalist methods such as Cellpose (4) and micro-SAM (5), yet this success has not translated to 3D, where segmentation remains challenging. A major bottleneck holding back generalist 3D segmentation models is a lack of labeled training data.

The scarcity of well-annotated 3D datasets comes from the laborious nature of manual labelling of 3D volumes (Fig. 1a). We illustrate this process in Movie S1, where an annotator goes through a stack slice-by-slice and roughly annotates one cell. The annotation of the cell is intentionally not pixel-precise, yet it still takes minutes to draw the labels in the x-y plane while keeping them consistent in the y-z and x-z planes. Labelling a single cell is laborious enough, so annotating hundreds of volumes with thousands of cells is nearly impossible. These difficulties prompted us to create a custom napari(6) plugin to simplify the manual annotation of masks with an ovoid shape (see Code and data availability). Indeed, without such a tool, even carefully labelled 3D volumes often show artefacts in z, where adjacent slices appear discontinuous along the x-z or y-z planes (Fig. 1b). Since the axial resolution of a microscope is typically lower than its lateral resolution, the upper and lower boundaries of a cell are hard to delineate, creating artefacts in manually labelled images (Fig. 1b). The scarcity of training data is also reflected in the tools themselves, such as StarDist (7) or Omnipose (8). Indeed, these network’s architectures can handle 3D volumes, but they do not make trained 3D models available, and the authors explicitly cite the lack of training data as being the bottleneck.

**Figure 1.**
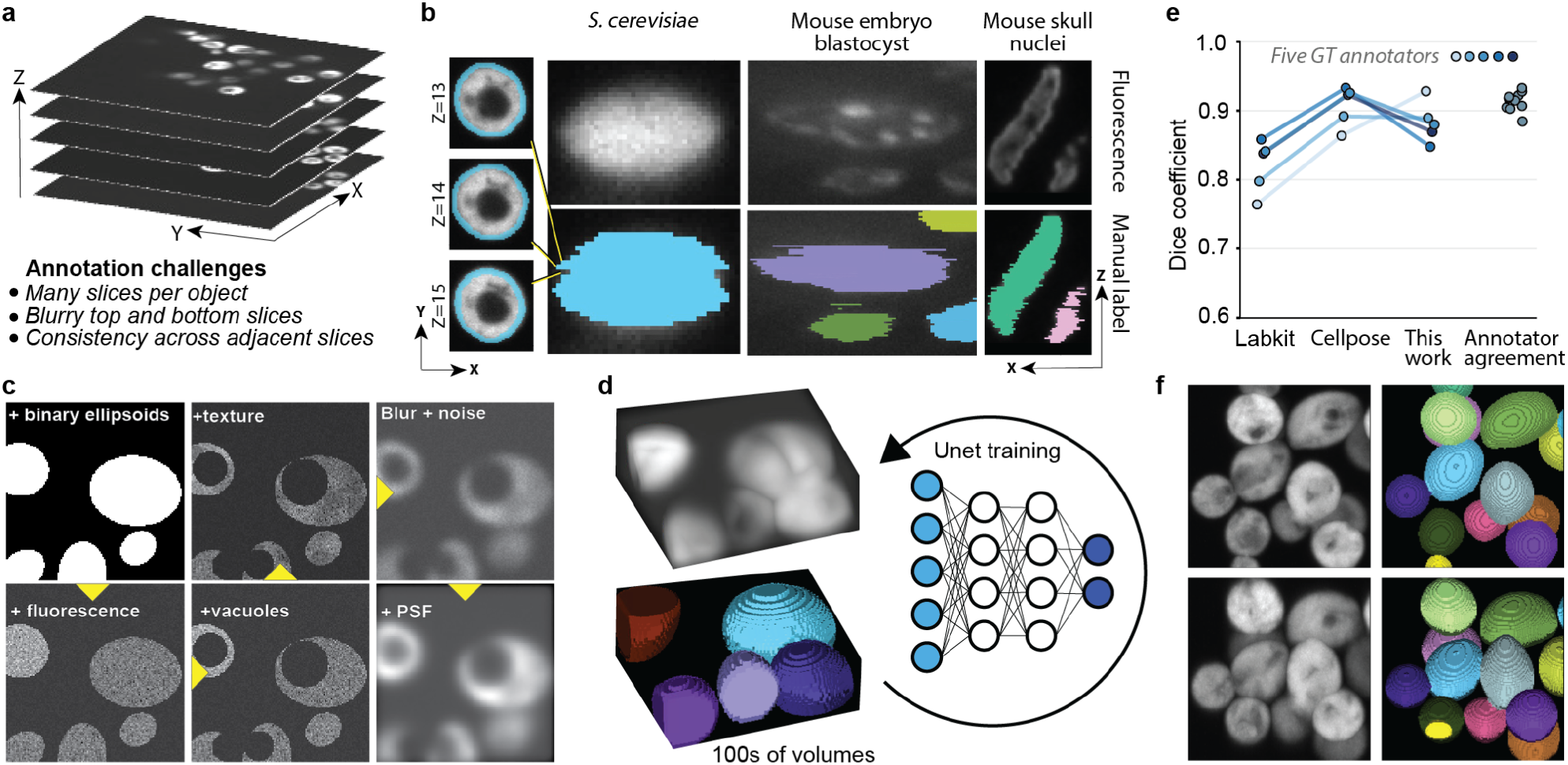
Synthetic volumes overcome training data scarcity for 3D cell segmentation. **a**, Manual annotation in 3D is laborious because each object spans many slices, because the uppermost and lowermost slices of each object are blurred by the axial point-spread function, and because every label should be continuous across adjacent slices. **b**, Obtaining annotations that are consistent across all (x, y, z) directions is particularly laborious. We annotated a yeast cell in x-y planes and show three of them, at z = 13, 14 and 15. Changing the view to a cross section along x-z reveals a discontinuity that should ideally be corrected, and such discontinuities are common in manually annotated data, e.g., in mouse embryo blastocysts (centre) (18), or mouse skull nuclei (right) (19). **c**, Strategy that we followed to generate synthetic volumes mirroring real 3D images of yeast cells. Binary ellipsoids are placed in a volume, assigned fluorescence intensities, given intracellular texture and vacuoles, degraded by blur and noise and finally convolved with a point-spread function. Yellow arrowheads indicate the order of steps. **d**, Because each volume is generated from its own mask, the image and its labels are obtained jointly, with pixel precision, and without effort. Hundreds of such pairs were used to train a 3D U-Net (15,20) with no real images and no manual annotation. **e**, Dice coefficient obtained on a real volume of yeast cells using Labkit (16), Cellpose (4) and the present method compared to the same volume annotated manually by five different persons. Lines join comparisons against the same manual annotation. The last column shows Dice values between pairs of manually annotated image masks. **f**, A real fluorescence volume of yeast cells (left) and the instance segmentation returned by our model (right); each colour denotes one cell. All images were visualized and rendered in napari (6).

Synthetic images offer a way around this bottleneck, because they provide labelled data without the need for manual annotation. Each volume is generated from a mask fixed in advance, so the cell boundaries are known exactly, and labelled datasets that would take months to trace by hand can be produced in minutes (Fig. 1c). Synthetic data has been used successfully to segment cells in two dimensions (9) and 3D approaches have been proposed, yet existing approaches are not completely independent of labelled training data (10,11), or do not reproduce the physical process of image formation, notably the point-spread-function (PSF) (12,13). Considering the PSF was key when simulating data of moving single particles (14) and we can anticipate that it is also important for trained networks to not over-segment into axially blurred regions.

Here we show that synthetic volumes generated with a minimal set of rules (Fig. 1c) are sufficient to train a standard 3D neural network (15) and reach state-of-the-art performance, provided that the volumes capture essential properties of real cells and of the imaging process, notably the presence of large vacuoles and the convolution of the data with a point-spread function. Our image generation workflow is designed in a modular way such that it can in principle be reproduced for other cell types.

The main steps of our generation workflow are illustrated in Fig. 1c. A volume of defined size, here 512 x 512 x 123 voxels, is first initialized at zero. Ellipsoids are then placed at random positions and orientations, with a new cell mask being accepted only if it does not overlap with existing ones beyond a set cut-off. Each cell is then assigned a fluorescence intensity drawn from a range, and an intracellular texture is added as small spheres of differing relative intensities. Dim and relatively large vacuoles are also placed within cells, and the volume is finally blurred, corrupted with multiplicative noise, and convolved with a pre-computed 3D kernel of a point-spread function. Finally, the volume can be used in full, cropped, or sub-sampled in Z to generate a training set with a given size and a fixed or varying anisotropies.

A model trained exclusively on these synthetic images (Fig. 1d), and therefore without any manual labelling, could segment real three-dimensional yeast volumes accurately (Fig. 1e). The model reached a mean Dice coefficient of 0.88, comparable with the state of the art represented by Cellpose (Dice = 0.90) and with the agreement between the five manual annotations of the same volume (mean Dice of 0.91, ranging from 0.88 to 0.92). A more naïve approach such as Labkit (16) reached an average Dice of 0.81 (Fig. 1e). This accuracy was not reached on our first attempt, and it came through iterations in which the required improvement was visible upon inspecting the predictions. A first model segmented cells accurately in the middle of each stack, but it failed at the uppermost and lowermost slices, irregularly extending masks beyond the cells and merging close neighbours. We reasoned that the cause was the point-spread function, which spreads fluorescence in real images along the optical z-axis more than within the x-y plane. Thus, we included this effect in the training set using first principles, by computing a theoretical point-spread function kernel with an established method (17) and convolving the synthetic volumes with it. A model trained on these new data no longer failed at the cell boundaries, but often left holes inside the cells, concentrated at the vacuole boundary. In real images the vacuole was indeed far darker than the cytoplasm, close to the contrast observable between cell foreground and background, so the model assigned those voxels to background. After we darkened the synthetic vacuole to the level of the background, the model no longer predicted holes within cells.

Our synthetic data is powerful for an additional reason: since each volume is built from its own mask, additional channels that carry information about cell geometry can be generated at no cost, to help the network learn to predict additional outputs or learn through auxiliary losses. Here, to solve the difficult task of instance segmentation we also generated a “centres volume” containing inner ellipsoids whose radii are 70% of those of each cell (34% of its volume). This channel can then be supervised together with the mask during training, so that at inference the centres are also predicted. The centres can then be used to seed a watershed and separate touching cells. Thus, from the mask and centres’ channels we can derive an accurate instance segmentation of yeast cells (Fig. 1f).

This simple strategy enabled us to produce instance segmentation masks of high-enough quality for unambiguous tracking or for measuring cell volumes, which we could not otherwise achieve with other approaches. This workflow can, in principle, be generalized to other cell types or subcellular structures and thus offers a general route to overcoming the annotation scarcity of labelled 3D data.

In light of these results, we propose that the future of 3D segmentation may rely less on new models than on methods to generate synthetic data that reproduce essential properties of diverse cells or structures of interest, which can be identified with a few iterations of model training and error inspection. Ultimately, generalist 3D segmentation models could be trained on synthetic data aggregated from numerous research laboratories.

## Methods

### Image acquisition

Volumes of *Saccharomyces cerevisiae* expressing a fluorescent cytoplasmic reporter were acquired on an Olympus IX83 microscope equipped with a Yokogawa CSU-W1 spinning-disc confocal scanner and dual Prime BSI sCMOS cameras, using a 60x/1.4 numerical aperture oil-immersion objective (108 nm lateral pixel size). Z-stacks (53 slices) were acquired at 0.25 µm steps.

### Synthetic data generation

Cell masks were produced by placing ellipsoids at random positions and orientations in a 512 x 512 x 123 voxel volume, accepting a new cell only when its overlap with existing cells stayed below 10% of either cell’s volume. Long-axis radii were drawn from a normal distribution (mean 30, 40 or 50 voxels depending on the volume, s.d. 3 voxels) and short-to-long axis ratios uniformly between 0.6 and 0.9. A concentric inner ellipsoid, with radii scaled to 70% of those of each cell, defined the centre label used later for instance seeding. Fluorescence images were then generated from these masks: background and per-cell mean intensities were drawn from distributions estimated from real images, and intracellular intensity texture was added as small spheres of varying relative intensity. A vacuole was placed in each cell and overwritten with background-level intensity. Volumes were degraded by Gaussian blur (**σ** = 3 voxels) and multiplicative noise (uniform, ±10%), down-sampled along z from 123 to 53 slices to match the axial spacing of the real acquisitions, and finally convolved with the point-spread function described below. A total of 504 volumes were generated.

### Point-spread function

A theoretical point-spread function was generated in Fiji using the PSF Generator plugin (17), based on the Born and Wolf 3D optical diffraction model, with a numerical aperture of 1.4, an emission wavelength of 520 nm, and a refractive index of 1.5. The resulting kernel was generated at 64 x 64 x 16 voxels (x, y, z) and normalized to unit sum. This single kernel was applied to every synthetic volume, after sub-sampling along z and before training, by fast Fourier transform convolution.

### Network training

We used the standard 3d_fullres configuration of nnU-Net v2 (15) with a custom trainer that predicts three independent binary output channels (cell mask, cell-cell overlap regions and cell centers), each supervised separately under a combined Dice and cross-entropy loss with deep supervision. The model was trained with a patch size of 28 x 256 x 256 voxels (z, y, x), with 1 x 3 x 3 kernels in the first stage and 3 x 3 x 3 kernels elsewhere, a batch size of 2 and the default nnU-Net optimizer, with the learning-rate schedule set for 5000 epochs. The model was trained on the 504 synthetic volumes alone (fold 0 of the default nnU-Net five-fold split), with no real images and no manual annotation. Training was stopped after 1800 epochs and the checkpoint saved at epoch 1200 was used for inference on real data as it minimized holes inside of predicted cell masks.

### Instance segmentation

Semantic predictions were converted to instances by seeded watershed, while accounting for the axial anisotropy. Connected components of the predicted centre channel were labelled and used as seeds for a watershed on the inverted distance transform of the predicted cell mask (after filling interior holes), constrained to foreground voxels so that background was never assigned a cell label.

### Evaluation

Segmentation accuracy on a real yeast volume was quantified as the Dice coefficient against each of five manual annotations of that volume, each containing about 45 cells (depending on the inclusion of very small buds and of partial cells at the edges). Manual annotation was aided by a custom napari plugin we developed for this purpose, which fits an ovoid from manually placed boundary points. Cellpose(4) and Labkit(16) were applied to the same volume for comparison; in Fig. 1e each line joins the three measurements obtained against one manually annotated volume. Labkit segmentations were obtained after marking foreground and background regions manually in the graphical user interface on the same volume. Cellpose was run with the pretrained Cellpose-SAM model and parameters matching the data (anisotropy 2.31, diameter 37 pixels).

## Code and data availability

Code for synthetic data generation, network training, instance segmentation and manual annotation using the napari plugin is available at https://github.com/SamhitaRR/synthetic-data-.

## Supplementary material

**Fig. S1.**
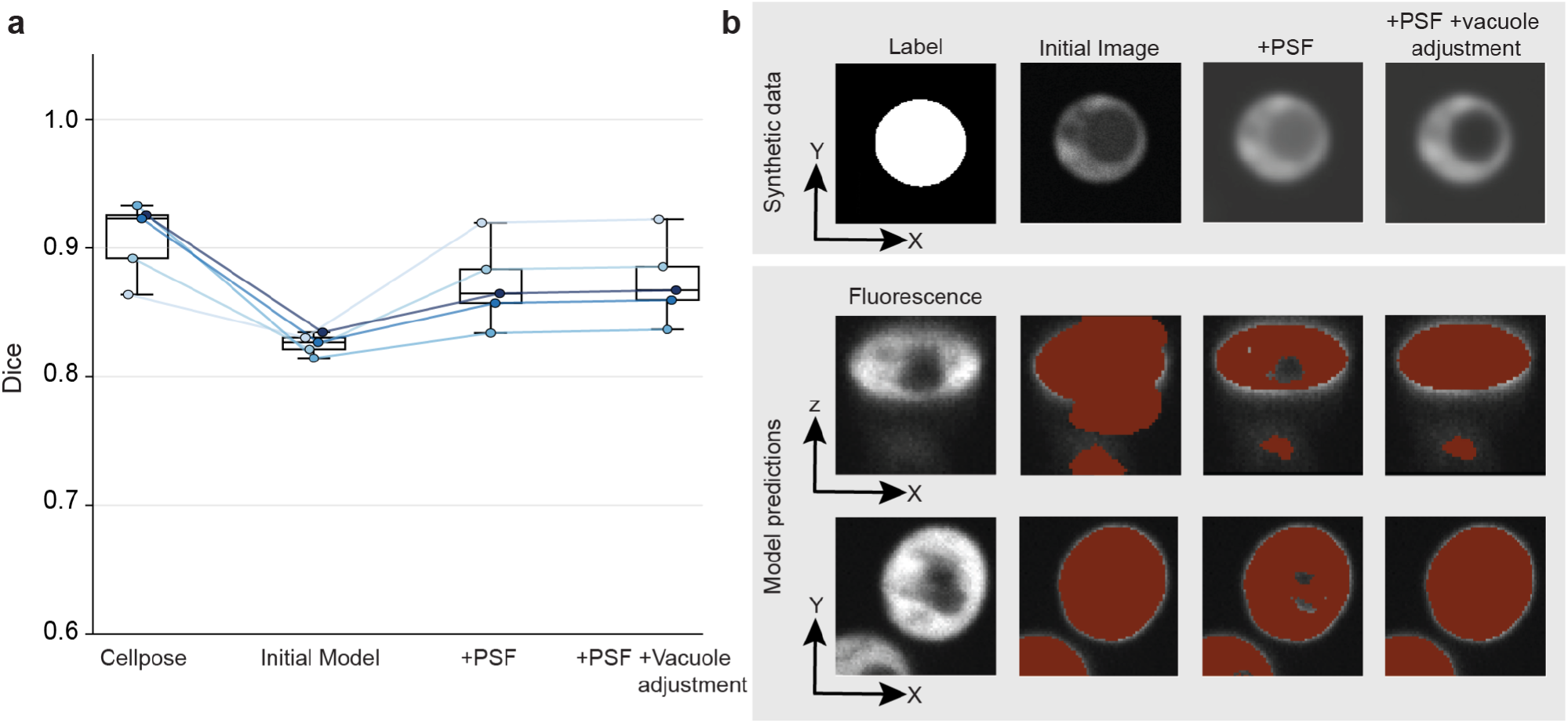
Convolving synthetic volumes with a PSF kernel enhances generalization to real data. (**a**) Dice coefficient on the real volume of Fig. 1e for models trained on three versions of the synthetic data: without convolution, with convolution with a kernel encoding a theoretical point-spread function (PSF), and with convolution and vacuoles set to background intensity. The latter is the model reported in Fig. 1. Each point is the Dice coefficient against one of the five manual annotations of Fig. 1e, lines join the same annotation, boxes show the median and interquartile range, and Cellpose is shown for reference. (**b**) Top, a synthetic cell (label and image) without and with application of the PSF, and after vacuole adjustment. Bottom, a real cell (fluorescence, x-z and x-y views) with the predictions of the three corresponding models in red. The convolution of the synthetic data with a PSF helps the model to not over-segment axially blurred regions in real images. The vacuole intensity adjustment helps the model to not leave holes inside cells, although this adjustment has little effect on the Dice value because holes are small relative to the cell size.

**Fig. S2.**
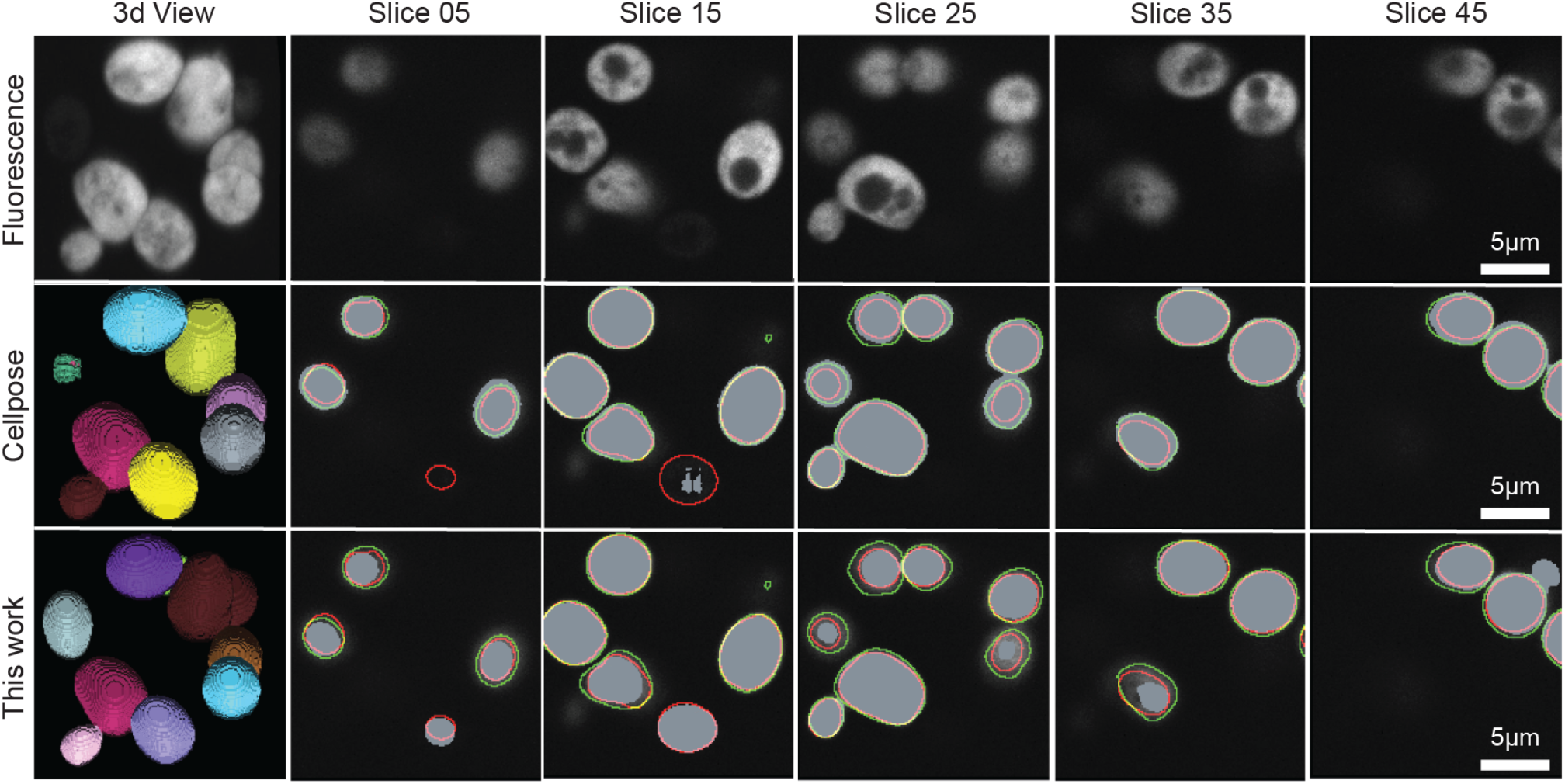
Illustration of predictions by Cellpose and our model on the same patch of real cells. The top row shows fluorescence data viewed in 3D with napari(6) (first column) and five corresponding slices sampled across the volume. The next rows show the instances (first column) and masks (subsequent columns) predicted by Cellpose(4) or by our model. Predicted masks appear in grey and the outlines show the manual annotations of two of the five annotators of Fig. 1e: the one drawing the smallest regions in red and the one drawing the most inclusive regions in green. Cellpose and the inclusive annotator both missed a dim cell at the bottom of the field. Except for this cell, the masks predicted by Cellpose tend to be larger than those of our model, which under-segments some cells (e.g., the lower cell in slice 35). Scale bars, 5 µm.

